# Recent African gene flow responsible for excess of old rare genetic variation in Great Britain

**DOI:** 10.1101/190066

**Authors:** Alexander Platt, Jody Hey

## Abstract

Population genomic studies can reveal the allele frequencies at millions of SNPs, with the numbers of observed low frequency SNPs increasing as more genomes are sequenced. Rare alleles tend to be younger than common alleles and are especially useful for studying demographic history, selection and heritability^1,2^. However, allele frequency can be a poor proxy for allele age, as genetic drift and natural selection can lead to alleles that are both rare and old. In order to allow joint assessments of allele frequency and allele age, a new estimator of allele age was developed that can be applied to variants of the lowest observed frequencies (singletons). By examining the geographic and age distribution of very rare variants in a large genomic sample from the UK^3^, we identify new evidence of gene flow from Africa into the ancestors of the modern UK population. A substantial proportion of variants with observed frequencies as low as 1.4 × 10^‒4^ are orders of magnitude older than can be explained without African gene flow and are found at much higher frequencies within modern African populations. We estimate that African populations contributed approximately 1.2% of the UK gene pool and did so approximately 400 years ago. These findings are relevant both to our understanding of human history and to the nature of rare variation segregating within populations: a variant that is rare because it is a recent mutation in the direct ancestor of the population will have had a very different evolutionary history than an ancient one that has persisted at high frequencies in a diverged population and only recently arrived through migration.

## Introduction

The two existing classes of methods for estimating the ages of genetic variants cannot be applied effectively to very rare variants. Methods that map allele frequencies to allele ages^4‒6^ cannot meaningfully distinguish among alleles that are already rare and greatly limiting the information they can convey. Other methods use the length of a shared haplotype, (or the rate of decay of linkage disequilibrium) among chromosomes that carry the rare allele to estimate the time since the carriers last shared a common ancestor^7,8^. However, these methods require that an allele occur at least twice in a sample. We developed an age estimator that can be applied to variants of any frequency including those that are observed just once. Instead of estimating the time since the common ancestor of all carriers of a variant, we estimate the time, *t*_*c*_, since the carriers of the variant last shared a common ancestor with a haplotype that does not carry the variant of interest. This time estimate serves as both an upper bound on the age of the variant itself as well as a measure of how long its lineage has been distinct from others in the sample. Being able to infer the age composition of very rare alleles allows us to unlock a new picture of how the forces shaping a gene pool may have recently changed, including recent shifts in ancestral composition.

We applied our method to estimate *t*_*c*_ for 21,992,410 of the rarest variants in the UK10k^3^ whole-genome population sequencing sample that has been filtered to remove close relatives and individuals of non-European ancestry. The distribution of estimated values revealed a dramatic excess of variation that is both old and rare ‐‐ well beyond what is predicted by previous models of UK or European human history. Figure 1 shows the means and standard deviations of the distributions of log(*t*_*c*_) values for variants found 2, 3, 4, 5,10, and 25 times in the UK10k sample of 3,621 individuals (7,242 haplotypes), and compares them with predictions from five published models of UK and European demographic histories^9‒13^ as well as new models with additional admixture events from an African population or diverged archaic human group. For all of the lowest frequency classes, the observed data contain variants that are far too old to have been generated by the published models (all of which returned mean simulated *t*_*c*_ distributions considerably smaller than for the observed data). The models proposed by Gutenkunst *et al*.^9^ and Gravel *et al*.^10^, the two published models with migration between Africa and Europe, return standard deviations similar to that of the observed data but with insufficient old alleles to substantially raise the predicted mean. For variants as common as those found 25 times (a frequency of ~0.35%), all models fit the observed distribution reasonably well.

**Figure 1.**
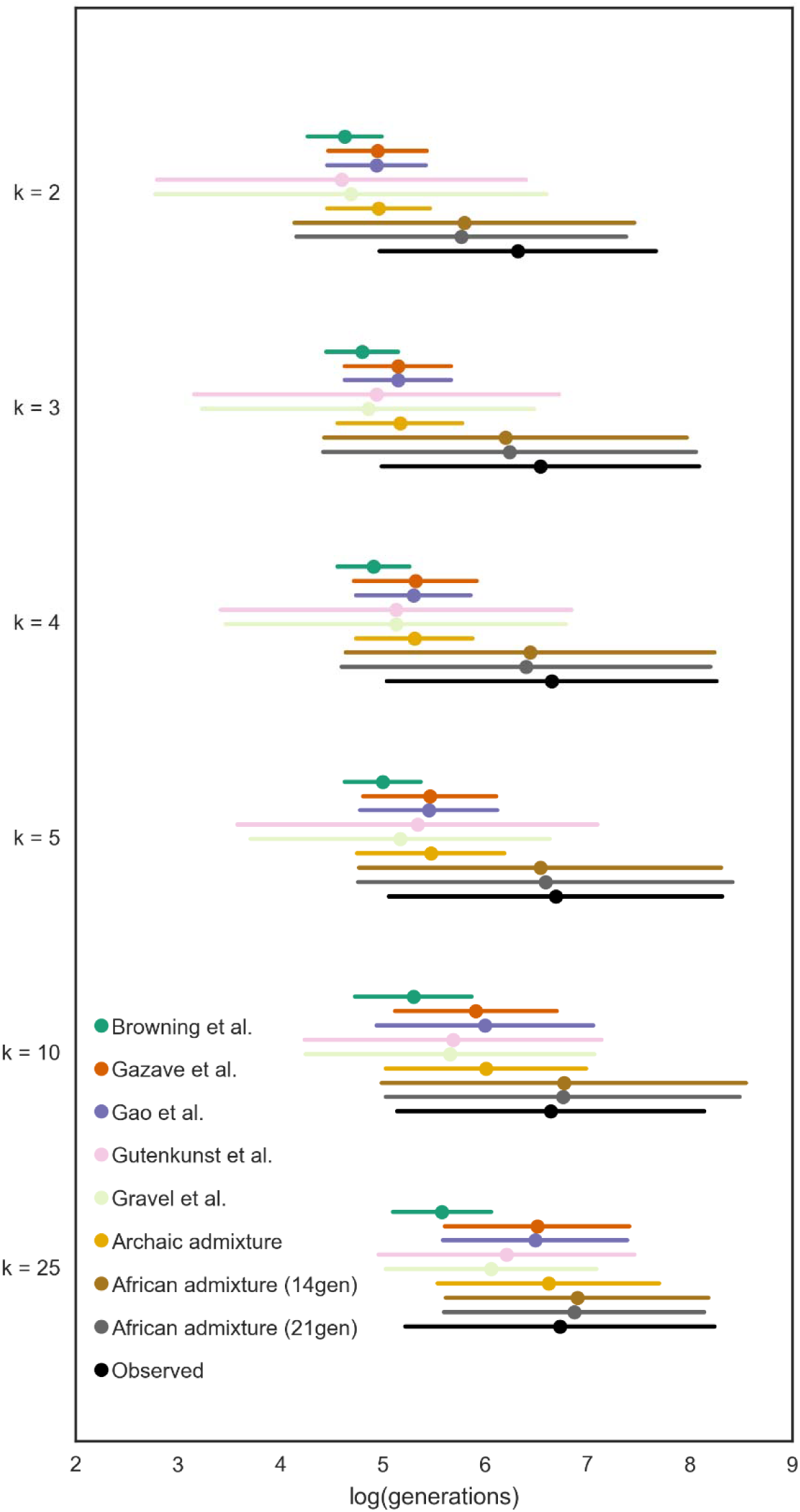
Distributions of log(*t*_*c*_) values. Variants of different frequencies in the UK10k data (k values) represented as mean ± one standard deviation of the log-transformed *t*_*c*_ values. The observed distribution is marked in black with other colors indicating expectations under proposed demographic histories of Britain and Europe.

Admixture from archaic humans will have introduced old alleles, and some of these are expected to appear at the lowest frequencies in the UK10K sample. However, we found that admixture with archaic humans does not introduce sufficiently rare alleles in the numbers necessary to explain the discrepancy. While the alleles introduced by such admixture are old, few of the introduced alleles end up in the relevant frequency classes at the time of sampling. When we turned to extant human populations as potential sources of old, very low frequency variants, we found an excellent fit with models that include recent admixture from African populations.

Figure 2 shows the fit of a series of models based on that by Gazave et al.^11^ with the addition of a recent migration event from an un-sampled African population to an ancestral UK population from which it separated 2,000 generations ago. The best fit model is one with 1.2% admixture 21 generations ago. This is a model in which ~10% of rare UK10k variants predate the human expansion out of Africa (see Extended Figure 2); have been segregating at moderate frequency within Africa; and were recently introduced to the UK population from Africa through migration.

**Figure 2.**
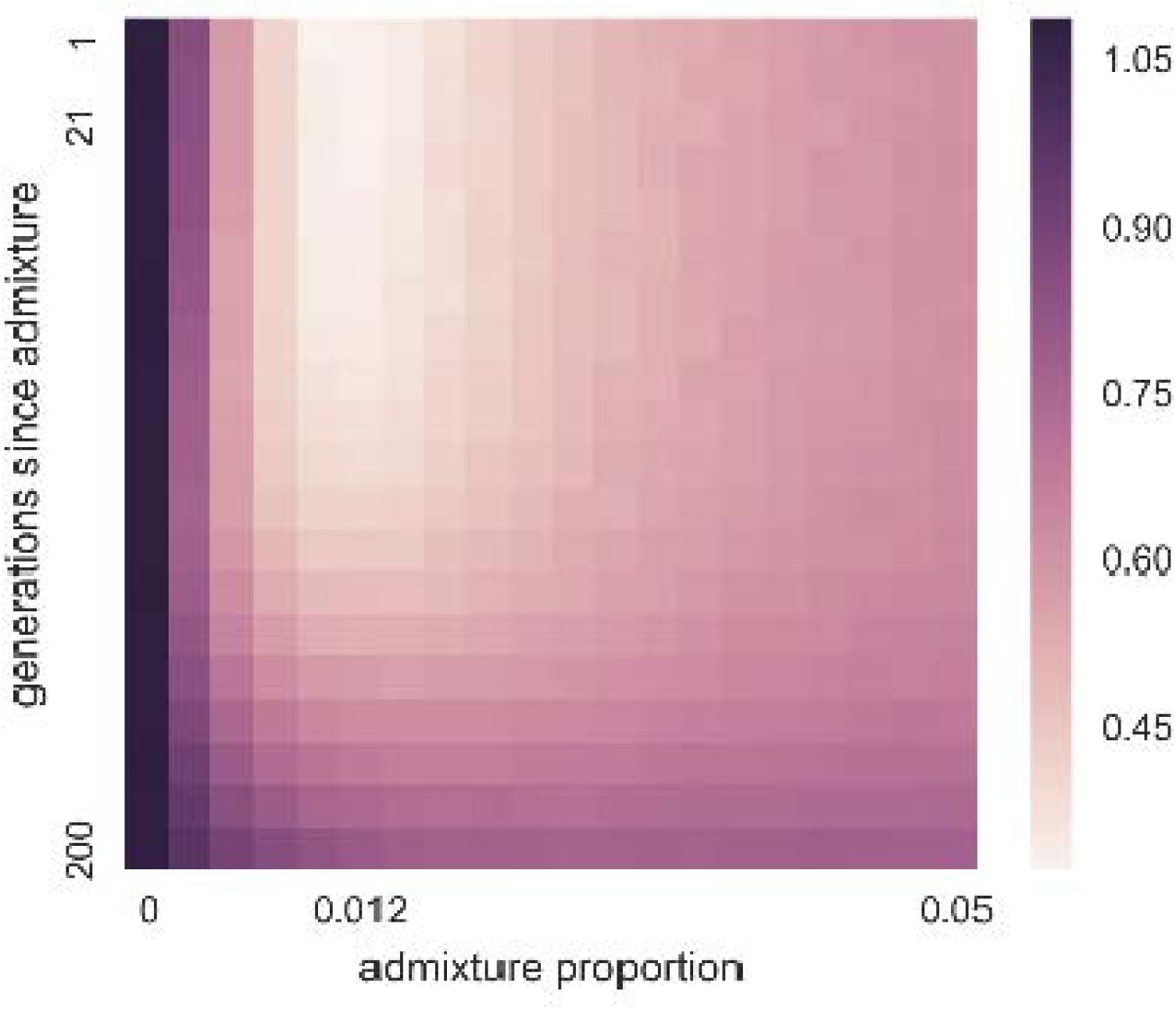
African admixture parameter optimization. Root mean squared log error between observed *t*_*c*_ values and those simulated over a grid of proposed values for timing and magnitude of African gene flow into the ancestors of the UK10k population.

If migration from Africa is responsible for substantially altering the age distribution of rare UK alleles we expect to find a substantial proportion of these rare British alleles at higher frequency in African populations than they are in the UK (or other European populations). We assessed all of the UK10k singleton variants, and variants found 25 times, for their presence and frequency within the populations of the 1000 genomes project phase 3 dataset^14^. Figure 3 shows that very rare UK10K alleles are typically at their rarest in Great Britain and present in higher frequencies in African populations than elsewhere in the world. In contrast, variants observed 25 times within the UK10k data show a very different pattern, as they are rarely found outside of Europe, but are found at relatively high frequency when they do occur. This is the pattern expected of alleles that have primarily *not* been recently introduced by migration, but rather have persisted both inside and outside of Africa since their origins before the population divergence at the time of the out of Africa expansion.

**Figure 3.**
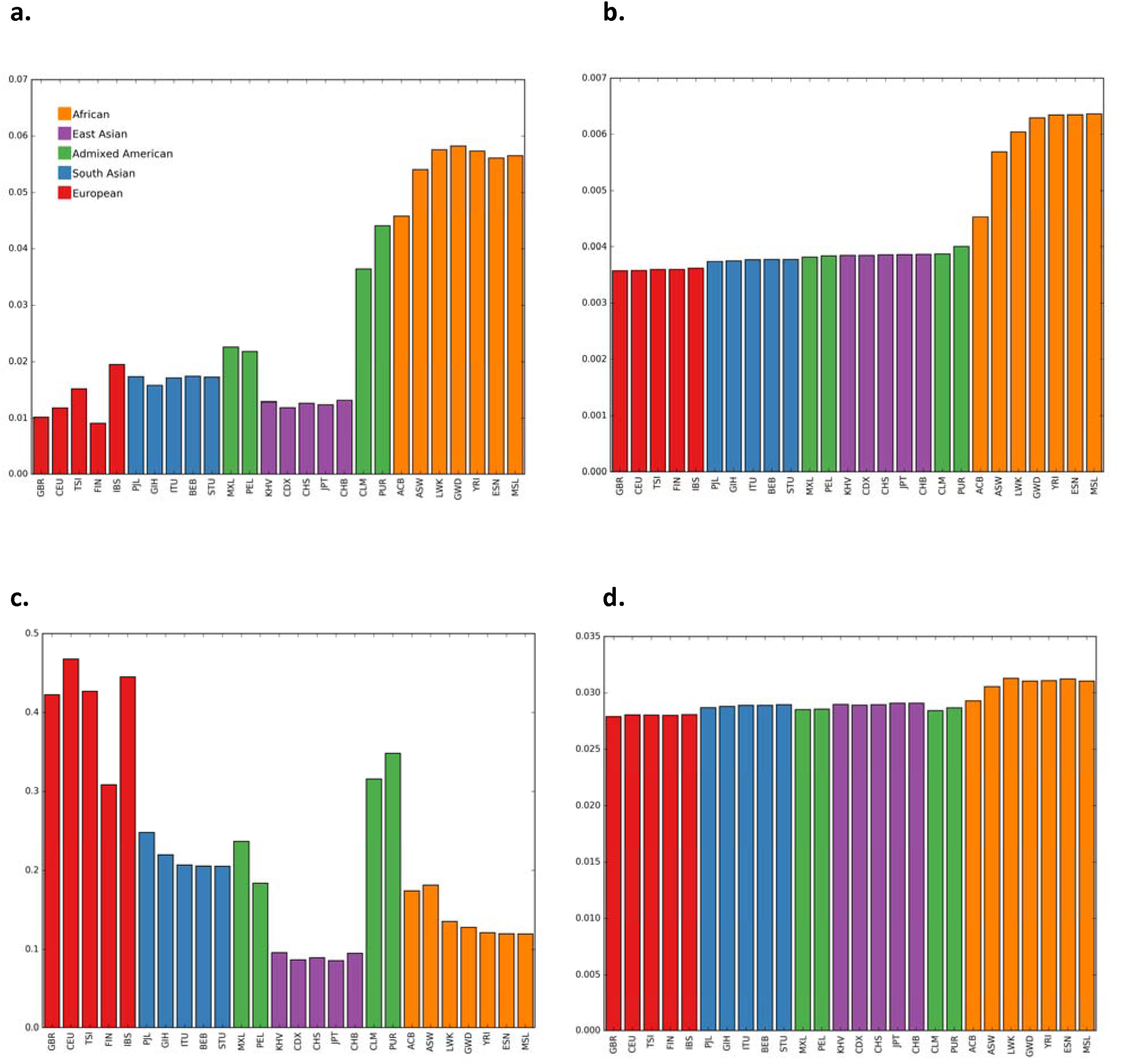
Geographic distribution of UK10k variants. Panels a. and b. reflect variants that are singletons in the UK10k data. Panels c. and d. reflect variants found 25 times in the UK10k data. Panels a. and c. describe the proportion of UK10k variants found in each population sample in 1000 Genomes Project data. Panels b. and d. describe the average frequency in the 1000 Genomes Project data of UK10k variants. 95% confidence intervals are all less than 0.4% of the bar heights for panels a. and b. and less than 4% of the bar heights for panels c. and d.

Extended data figure 1 shows that the distributions of *t*_*c*_values for UK10k variants that are found in African populations in the 1000 Genomes Project data are considerably older than those that are not. Similar results are found comparing the UK10K variants to those sequenced in the African samples of the Simons Diversity Panel^15^.

While figure 2 indicates support by the data for models with ^~^1.2% admixture, the overall distribution of rare allele ages is consistent with a range of models that vary in the timing of the admixture event. We further refined the estimate of the time of admixture by explicitly modelling the distribution of the numbers of introgressed alleles observed among the individuals in the UK10K sample. In a random-mating model with admixture occurring over a short period of time, the introgressed alleles will come to be spread fairly evenly in the population over a small number of generations. However, as shown in figure 4, the distribution of numbers of alleles, for alleles that occurred twice in the UK10K sample with *t*_*c*_ values greater than 2,500, shows a considerable clustering across individuals, with 13% of individuals harboring ^~^40% of all such variants. This clumping is suggestive of recent introgression, but could also be consistent with older introgression with the decay of clustering slowed by nonrandom mating. As shown in figure 4c, the observed distribution is well fit by a broad range of models of assortative mating, all implying a time of admixture 11-14 generations before sampling. The model ‘African admixture (14gen)’ in figure 1 represents the expected distribution of log(*t*_*c*_) values for a 1.2% admixture event 14 generations before sampling.

**Figure 4.**
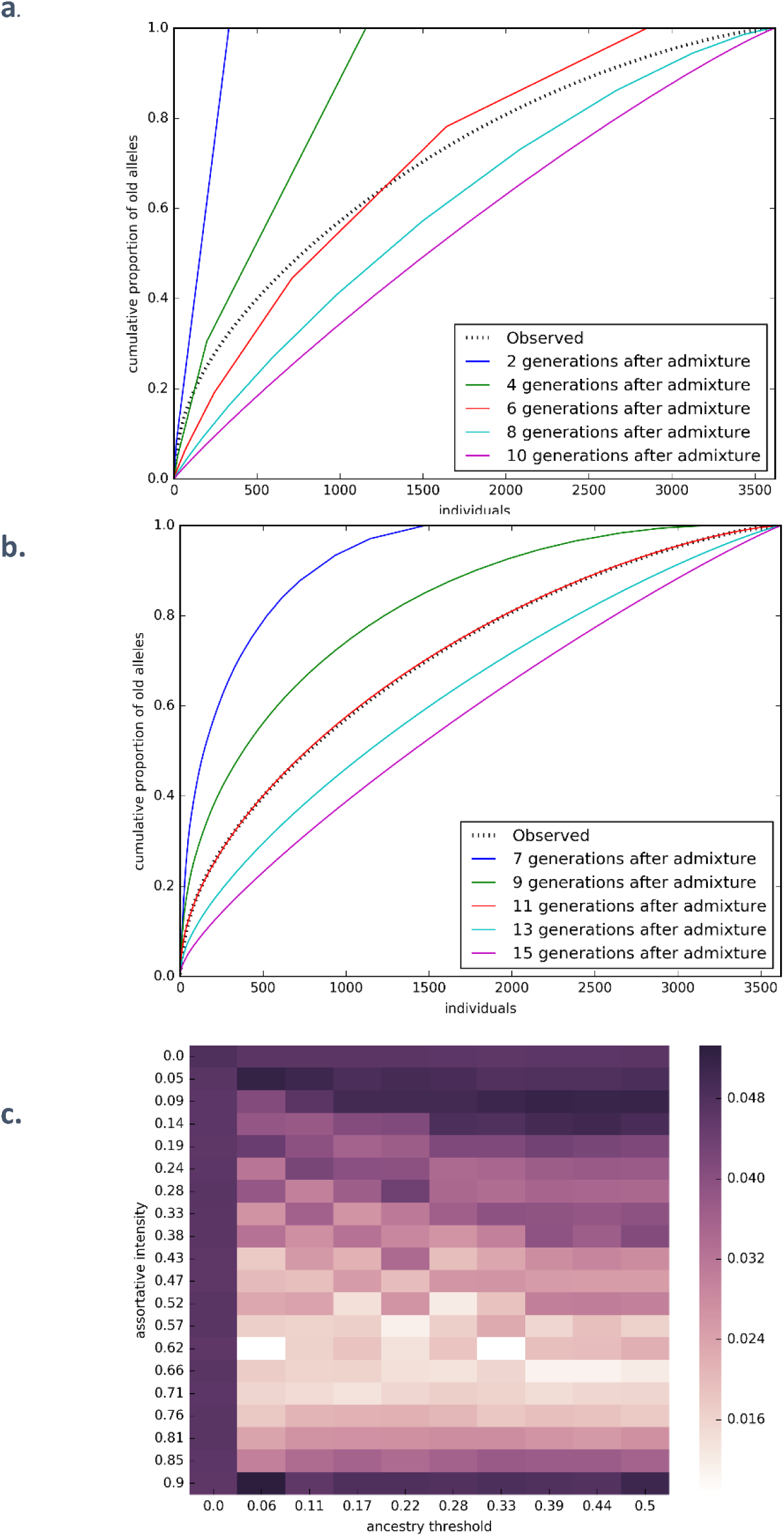
Cumulative proportion of doubleton alleles more than 2,500 generations old carried by individuals ordered from most to fewest old doubleton variants. Observed distribution is marked in dotted black. Colored lines are simulations of 1.2% admixture at different historic time points and assuming (a) random mating or (b) assortative mating with a threshold of 22% and intensity of 63%. Panel (c) shows the minimum (across ages) root mean square error between the observed cumulative proportion of old doubletons and that predicted under a range of models of non-random mating. Dark bars at the top and left indicate poor fit of random mating models. Light bar across the middle indicates similar support for models with a wide range of ancestry thresholds.

With African gene flow modeled as a point admixture event, we estimate 1.2% of the ancestors of the UK10k population came from Africa 11-14 generations ago. A generation time of 29 years^16^ places the migration event early in the era of the “First” British Empire when Britain was actively establishing colonies in West Africa and the West Indies^17^. During this period there was a notable rise in the Black British population, often as household attendants to returning sea captains and colonists or as former slaves from Spain and Portugal^18,19^.

This model of demographic influx is one in which many alleles that arose before the expansion out of Africa rose to (or remained at) relatively high frequencies within a much larger African population while being lost or excluded from the smaller ancestral European population. These alleles then had the opportunity to be reintroduced to the European population in very small quantities through recent migration. We are not excluding the possibility that the gene flow that shifted the distribution of UK10k *t*_*c*_ values was a more complicated phenomenon than a single migration event. With relatively higher levels of African gene flow into Southern European populations^20,21^ it is probable that some proportion of the African variants we find in the UK10k population did not come by immigration *directly* from Africa but by more circuitous means.

It has been argued that rare variants should fall almost exclusively into the class of recent mutations, and that rare variants are unlikely to be found outside of their population of origin^10,22^. This conclusion was drawn from a model fit from a smaller sample where the rarest variants were at considerably higher population frequencies than the rarest variants in the UK10k data. As shown in Figure 2, as a variant rises in frequency even to 0.0035 frequency (k=25 in the UK10k sample), the probability of finding it outside of Europe drops dramatically. It is predominantly among the very rarest of variants that we see the substantial impact of recent migrants.

## Methods

### UK10k population data

We use the mapping sample of 3,621 individuals from the UK10k data set that has been filtered by the UK10k consortium to remove close relatives and individuals of recent non-European ancestry. These include genomes from the ALSPAC cohort, which focused on the Avon region, and the TWINSUK cohort which includes samples from across the UK. We masked all CpG / TpG transversion polymorphisms to avoid homoplasy and reduce heterogeneity in the mutation rate. Haplotype phase for variants found two or more times was inferred by the UK10k consortium using SHAPEIT^23^. Haplotype phase for singleton variants was assigned to maximize their *t*_*c*_ values as branches of gene trees that harbor mutations are expected to be longer than those that do not. The *t*_*c*_ values for singleton variants were not used in fitting the demographic models.

### Estimating *t*_*c*_ values

We estimate the *t*_*c*_ value of a singleton variant as a function of the length of the maximum shared haplotype (*msh*) that extends from it in either direction. Starting at the site of the variant, we find the pairwise alignment with the rest of the sample that has the longest perfect match before the first discrepancy. This is done independently in the 5’ and 3’ directions giving a pair of *msh* observations that need not arise from the same alignment. We assume that the alignments that generate the *msh*i values are between the variant-carrying haplotype and its closest relatives. We model each *msh* value as the distance to the closest mutation or recombination event to have occurred on either the external branch leading to the singleton variant (which create discrepancies between the variant-carrying haplotype and all other haplotypes in the sample) or on its first sister branch (which create discrepancies between the variant-carrying haplotype and all of its closest relatives). Each *msh* is then treated as the distance to the closest event in a Poisson process with a density parameter equal to the sum of the mutation and recombination rates times the sum of two branch lengths: the external branch along which the singleton variant arose, which has length *t*_*c*_, and its sister branch, which has some length 0. With a uniform per-base recombination rate *ρ* and mutation rate *μ*, the probability density is an exponential distribution:

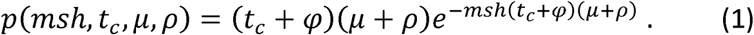

In practice we relax the assumption of uniform recombination rate and introduce two modifications to how we record the *msh* values: first, to minimize artifacts from mis-phased singleton variants, we do not end a maximum shared haplotype tract if a discrepancy is due to a singleton variant. Second, while our derivation models *msh* as stopping at a recombination event, such events cannot be directly observed, and so the measured *msh* necessarily continues until a (non-singleton) mutation is encountered that is not shared between the aligned haplotypes. This is expected to occur very shortly after a recombination event as nonshared mutations accrue at a much higher rate beyond the first recombination event.

The probability density of the length of the sister branch, conditional on *t*_*c*_, is derived from a hazard model of the rate of coalescent forward in time^24^.

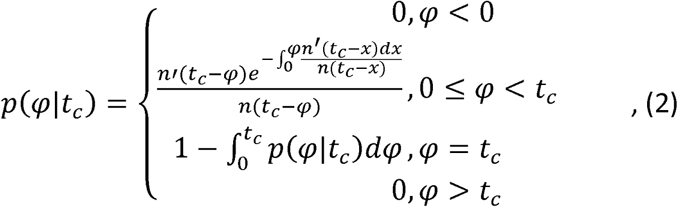

where *n*(*x*) is the number of ancestral of lineages ancestral to sampled individuals that existed *x* generations in the past. Using an approximation for *n*(*x*)^25^ and changing variables, we get:

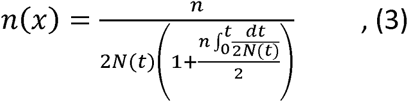

where *n* is the the number of lineages at time 0 (i.e. the sample size), and *N*(*t*) is the size of the population *t* generations in the past.

For a given SNP we record *χ* = *u*(*x*_5′_ + *x*_3′_) + *c*_5_, + *c*_3_, where *x*_5_, and *x*_3_, are the *msh* distances on the 5′ and 3′ sides of the focal base position, in base pairs, and *c*_5′_ and *c*_3′_ are the same distances in Morgans. Working in absolute recombination distances allows a relaxation of the assumption of uniform recombination rate. Then the likelihood can be written as

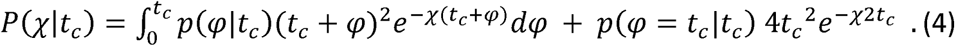

This expression is readily simplified for a constant or exponential growth population model, while more complex demographic histories may require numeric integration. In either case the maximum likelihood estimate of *t*_*c*_ can be found by numerical optimization.

Because the demographic model enters (4) through the density of *φ*, which makes a minority contribution to the likelihood, the choice of the form of the demographic model (i.e. constant versus exponential versus multi-phase) actually has very little impact on parameter estimates (see Extended Data Figure 3). This insensitivity of the estimator to demographic assumptions is a useful property of a statistic that is to be used to uncover components of demographic histories. We use a fixed *N* = 5,000 for all analyses of the UK10K variants.

For variants that are not singletons, we use a composite likelihood to estimate the value of *t*_*c*_ that best fits all of the *msh* values simultaneously. We approximate their joint likelihood as the product of the likelihoods of the *msh* values for each copy of the variant considered independently, each in turn being compared to the rest of the sample that do not carry the rare allele.

While statistics derived from maximum values of observed data (such as the maximum length of a shared haplotype in our estimator) are often poorly behaved with large biases and strong sensitivities to model misspecification or stochastic noise, our estimator performs quite well, even under adverse conditions, such as when using *msh* values from statistically phased haplotypes. We demonstrate this through coalescent simulations generated with msprime^26^. As shown in Extended Data Table 1 and Extended Data Figure 4, for all of our simulations the bias was low. Precision was good for simulated populations of constant size but was low when estimating *t*_*c*_ for very young singleton variants in models of extreme recent population growth. However even in these cases, the bias is small and precision may be sufficiently good to allow studies of many variants. For non-constant population size models precision returns to the higher levels found in constant-sized populations for variants of slightly higher frequencies.

### Evaluating demographic models

We estimate the mean and standard deviation for a given model by simulating 500 independent coalescent trees with sample sizes of 7,242 haplotypes using msprime. For a low frequency allele observed *k* times we identify all of the branches that are ancestral to *k* leaves and calculate the average of their starting ages weighted by the ratio of their lengths to the total length of branches with *k* leaves across the set of trees.

We evaluated the distributions of *t*_*c*_ values under eight different demographic scenarios. Browning et al.^13^ is a single-population model fit to UK10k data. Gazave et al.^11^ and Gao et al.^12^ are single-population models fit to the European-American population of the NHLBI Exome Sequencing Project^27^. Gutenkunst et al.^9^ and Gravel et al.^10^ are three-population models (African, European, and Asian) fit to data from the Environmental Genome Project^28^ and 1000 Genomes Project respectively, with all of the samples treated as coming from the European population. Archaic admixture is a model of ours that augments Gazave et al. with an archaic population that diverged 25,000 generations in the past and contributed 3% to the European population 2,000 generations ago. Our models of African admixture (African admixture (21gen) and African admixture (14gen)) use the Gazave et al. demographic model but add an additional 1.2% contribution of African lineages to the European population 21 or 14 generations ago respectively. Model fit is ascertained by calculating the root mean squared error of the means of the log (*t*_*c*_) values for variants occurring 2, 3, 4, 5,10, and 25 times in the UK10K data.

### Estimating migration parameters

Parameters describing the timing and quantity of African admixture were ascertained in a two-stage process. First, root mean squared log error of the mean *t*_*c*_ values for variants occurring 2, 3, 4, 5,10, and 25 times in the UK10K data were evaluated for each cell of a 20 x 20 twodimensional grid of parameter values, with admixture varying between 0 and 5% and the time of admixture varying from 1 to 200 generations prior to sampling.

Second, we refined our estimate of the time of admixture by modeling the spread of immigrant doubleton alleles through the population. At the time of admixture, all of the immigrant alleles will reside in a proportion of the population equal to the admixture proportion. In subsequent generations these alleles spread across more individuals. Using the admixture proportion of 1.2% identified in the previous grid search, we fit a time since admixture using a Wright-Fisher forward simulation with a two-parameter model of associative mating. A threshold parameter defines two ancestry groups and an intensity parameter describes the degree of preference for within-group mating. In each generation the population is divided into two ancestry groups: individuals with more than the threshold African ancestry and those with less. For each offspring in the next generation, one parent is drawn at random from the population. With probability equal to the intensity parameter the second parent is chosen at random from within the same ancestry group as the first parent. With probability one minus the intensity parameter the second parent is chosen at random from the entire population. Each offspring is assigned an ancestry proportion equal to the mean of its parents.

## Extended Data

### Tables

**Extended Data Table 1.** Performance characteristics of *t*_*c*_ estimator. Properties of estimated *t*_*c*_ values compared to true simulated *t*_*c*_ values, a) attributes of variants found *k* times in the sample, b) simulations of either constant-sized populations or one with recent exponential growth, c) haplotype phasing taken directly from the simulation (known) or statistically inferred from simulated diploid genotypes, d) root mean squared log error of estimator, e) average signed log error, f) Pearson’s *r* statistic.

| $k^a$ | demography<sup>b</sup> | phasing<sup>c</sup> | error<sup>d</sup> | bias<sup>e</sup> | correlation<sup>f</sup> |
| --- | --- | --- | --- | --- | --- |
| 1 | constant | Known | 0.16 | 0.09 | 0.87 |
| 5 | constant | Known | 0.11 | 0.04 | 0.84 |
| 10 | constant | Known | 0.10 | 0.02 | 0.82 |
| 25 | constant | Known | 0.11 | -0.02 | 0.69 |
| 1 | constant | inferred | 0.39 | -0.16 | 0.41 |
| 5 | constant | inferred | 0.11 | 0.03 | 0.82 |
| 10 | constant | inferred | 0.10 | 0.01 | 0.81 |
| 25 | constant | inferred | 0.11 | -0.03 | 0.68 |
| 1 | growth | Known | 0.17 | 0.11 | 0.55 |
| 5 | growth | Known | 0.10 | 0.06 | 0.76 |
| 10 | growth | Known | 0.09 | 0.04 | 0.85 |
| 25 | growth | Known | 0.09 | 0.01 | 0.81 |
| 1 | growth | inferred | 0.22 | -0.05 | 0.26 |
| 5 | growth | inferred | 0.10 | 0.03 | 0.70 |
| 10 | growth | inferred | 0.09 | 0.02 | 0.82 |
| 25 | growth | inferred | 0.09 | -0.01 | 0.80 |

### Figures

**Extended Data Figure 1.**
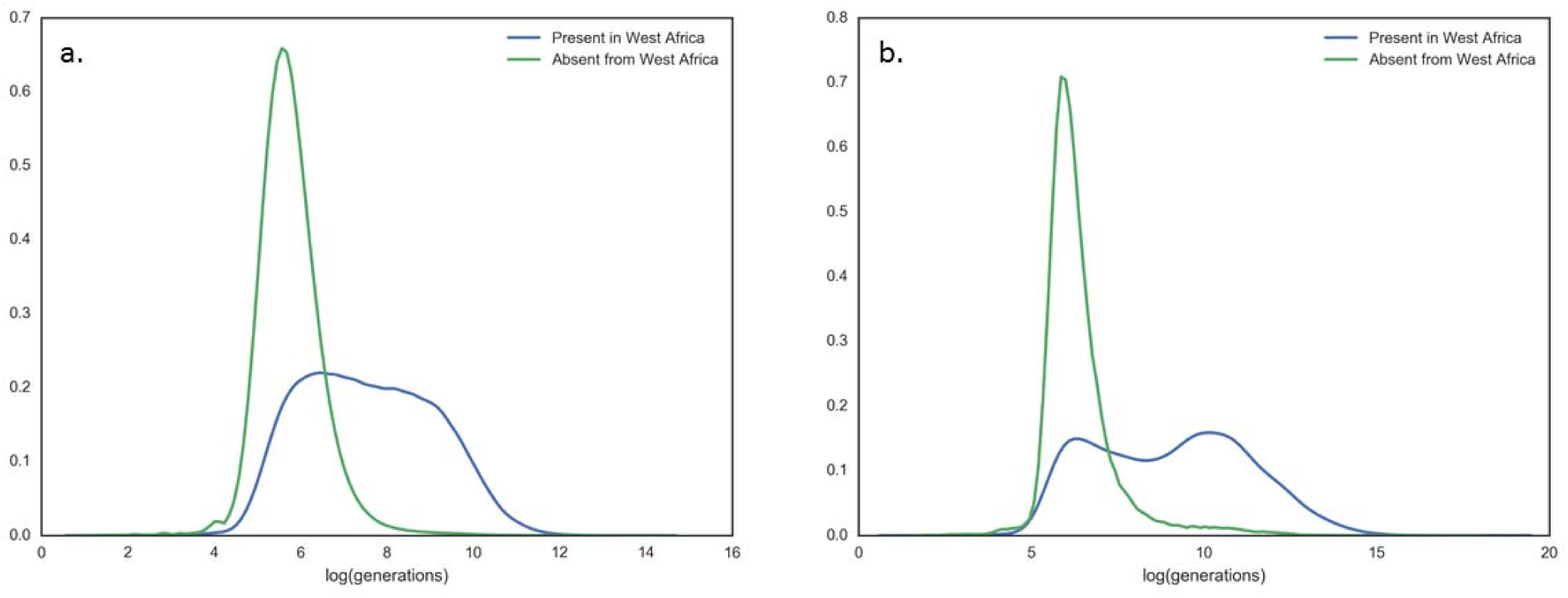
Age distribution of African variants. Distributions of *t*_*c*_ values for UK10k singleton variants (a) and variants found 25 times (b) grouped by presence or absence in any West African population of the 1000 Genomes Project.

**Extended Data Figure 2.**
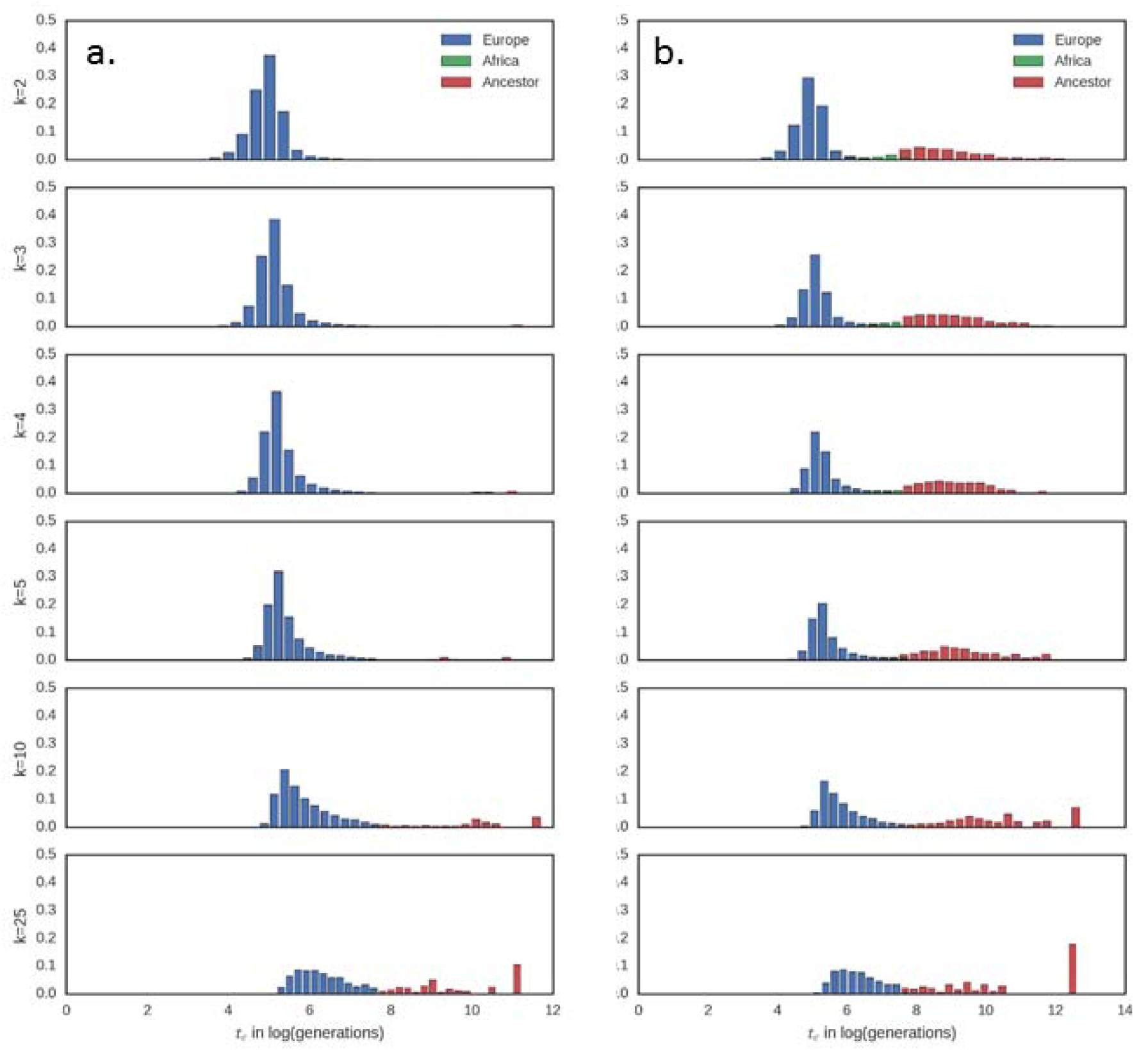
Expected distributions of *t*_*c*_ values by age and population. Simulation results for the expected distribution of *t*_*c*_ values colored by the population in which the coalescent event took place. Demographic models are (a) Gazave et al. and (b) African admixture (21gen).

**Extended Data Figure 3.**
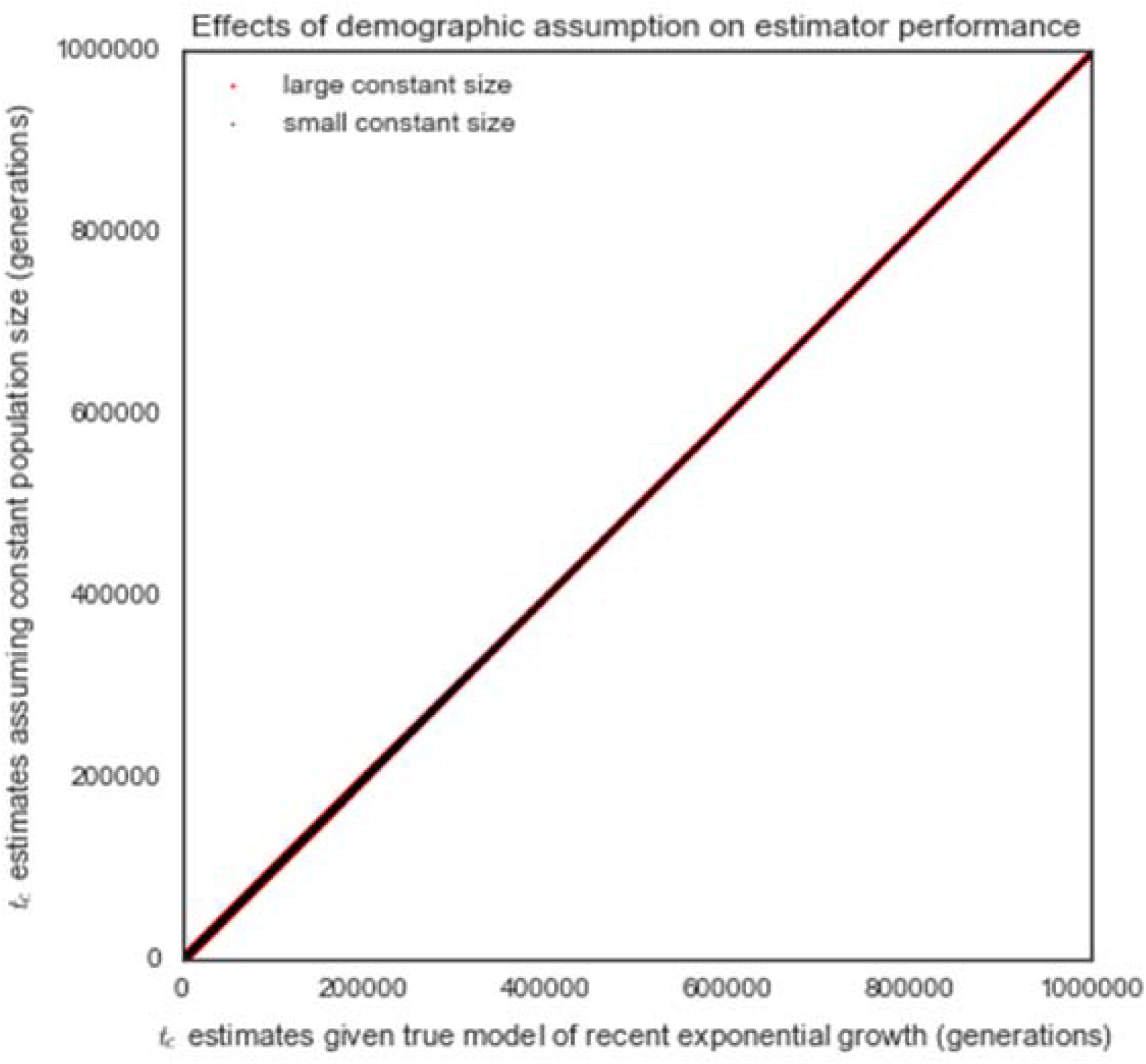
Estimator robustness to demographic model assumptions. Distribution of estimated *t*_*c*_ values for simulated singleton variants estimated assuming a constant-sized population of 5,000 (black) and 500,000 (red) as a function of the value estimated assuming the correct generative model of exponential population growth from 5,000 to 500,000 over the last 120 generations.

**Extended Data Figure 4.**
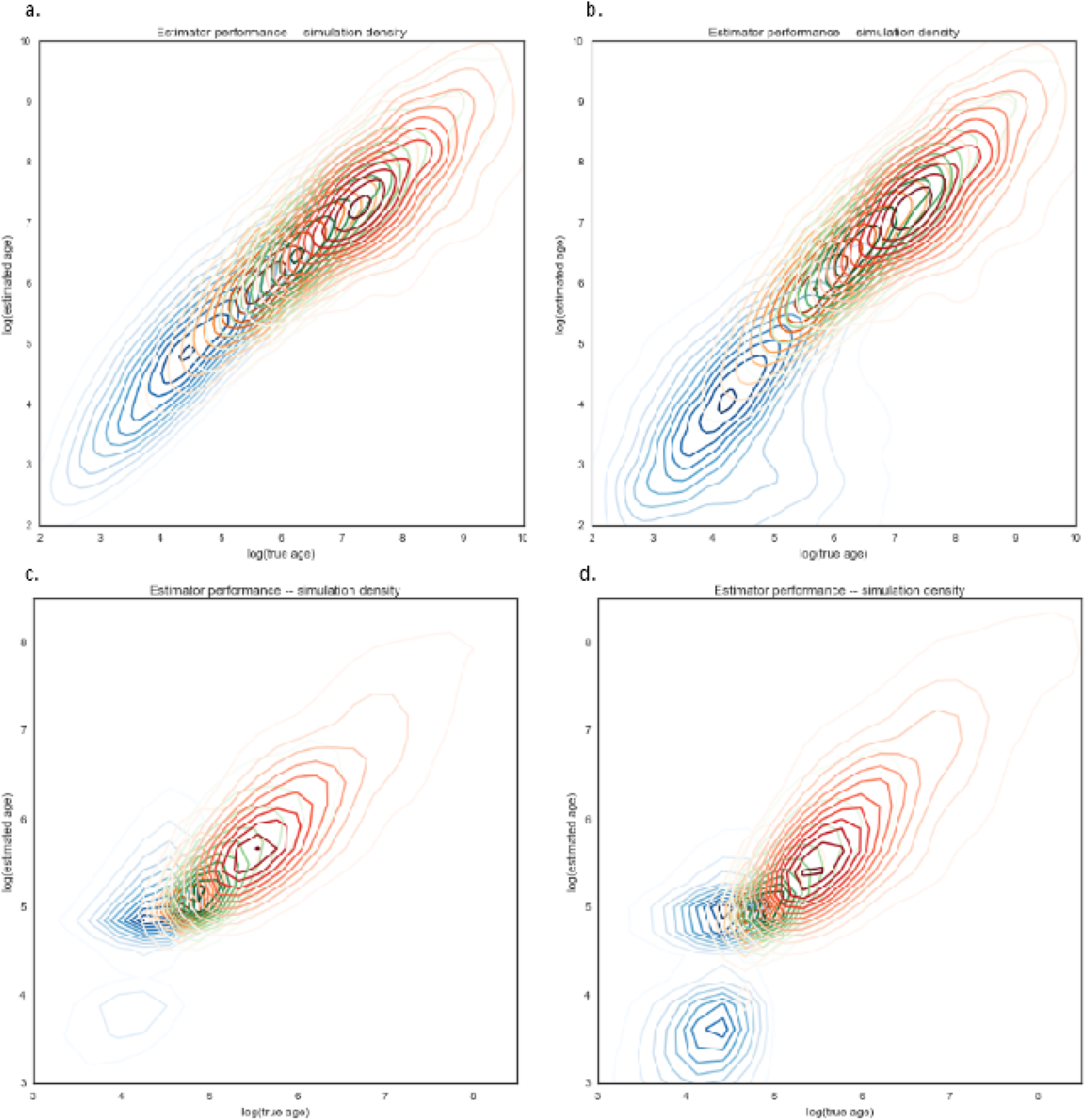
Estimator performance. Density plots for estimation of *t*_*c*_ values as a function of true *t*_*c*_ values in simulated data. Panels a. and b. are simulated constant-sized populations of 10,000. Panels c. and d. are populations that have grown exponentially from 5,000 to 50,000 over the last 120 generations. Panels a. and c. are evaluated on true known haplotypes. Panels b. and d. use statistically phased haplotypes from simulated genotypes. Densities of singleton variants is indicated in blue, variants sampled five times in orange, 10 in green, and 25 in red.

## Citations

1. Zuk, O. et al. Searching for missing heritability: designing rare variant association studies. Proc. Natl. Acad. Sci. 111, E455–E464 (2014).

2. Henn, B. M., Botigué, L. R., Bustamante, C. D., Clark, A. G. & Gravel, S. Estimating the mutation load in human genomes. Nat. Rev. Genet. 16, 333–343 (2015).

3. Consortium, U. & others. The UK10K project identifies rare variants in health and disease. Nature 526, 82–90 (2015).

4. Kimura, M. & Ohta, T. The Age of a Neutral Mutant Persisting in a Finite Population. Genetics 75, 199–212 (1973).

5. Griffiths, R. C. & Tavaré, S. The age of a mutation in a general coalescent tree. Commun. Stat. Stoch. Models 14, 273–295 (1998).

6. Fu, W. et al. Analysis of 6,515 exomes reveals the recent origin of most human protein-coding variants. Nature 493, 216–220 (2013).

7. Slatkin, M. & Rannala, B. Estimating the age of alleles by use of intraallelic variability. Am. J. Hum. Genet. 60, 447–458 (1997).

8. Mathieson, I. & McVean, G. Demography and the age of rare variants. PLoS Genet 10, e1004528 (2014).

9. Gutenkunst, R. N., Hernandez, R. D., Williamson, S. H. & Bustamante, C. D. Inferring the joint demographic history of multiple populations from multidimensional SNP frequency data. PLoS Genet 5, e1000695 (2009).

10. Gravel, S. et al. Demographic history and rare allele sharing among human populations. Proc. Natl. Acad. Sci. 108, 11983–11988 (2011).

11. Gazave, E. et al. Neutral genomic regions refine models of recent rapid human population growth. Proc. Natl. Acad. Sci. 111, 757–762 (2014).

12. Gao, F. & Keinan, A. Inference of Super-exponential Human Population Growth via Efficient Computation of the Site Frequency Spectrum for Generalized Models. Genetics 202, 235–245 (2016).

13. Browning, S. R. & Browning, B. L. Accurate Non-parametric Estimation of Recent Effective Population Size from Segments of Identity by Descent. Am. J. Hum. Genet. 97, 404–418 (2015).

14. The 1000 Genomes Project Consortium. A global reference for human genetic variation. Nature 526, 68–74 (2015).

15. Mallick, S. et al. The Simons Genome Diversity Project: 300 genomes from 142 diverse populations. Nature 538, 201–206 (2016).

16. Fenner, J. N. Cross-cultural estimation of the human generation interval for use in genetics-based population divergence studies. Am. J. Phys. Anthropol. 128, 415–423 (2005).

17. Canny, N., Canny, N. P. & Low, A. The Oxford History of the British Empire: Volume I: The Origins of Empire. (OUP Oxford, 2001).

18. Shyllon, F. O. Black People in Britain 1555-1833. (Institute of Race Relations, 1977).

19. Kaufmann, M. Africans in Britain: 1500-1640. (Thesis DPhil-University of Oxford, 2011).

20. Moorjani, P. et al. The History of African Gene Flow into Southern Europeans, Levantines, and Jews. PLOS Genet. 7, e1001373 (2011).

21. Botigué, L. R. et al. Gene flow from North Africa contributes to differential human genetic diversity in southern Europe. Proc. Natl. Acad. Sci. 110, 11791–11796 (2013).

22. Nelson, M. R. et al. An abundance of rare functional variants in 202 drug target genes sequenced in 14,002 people. Science 337, 100–104 (2012).

23. O’Connell, J. et al. Haplotype estimation for biobank-scale data sets. Nat. Genet. 48, 817–820 (2016).

24. Platt, A. Time-conditional properties of branches in coalescent gene trees. bioRxiv 189209 (2017). doi:10.1101/189209

25. Slatkin, M. & Rannala, B. Estimating the age of alleles by use of intraallelic variability. Am. J. Hum. Genet. 60, 447 (1997).

26. Kelleher, J., Etheridge, A. M. & McVean, G. Efficient Coalescent Simulation and Genealogical Analysis for Large Sample Sizes. PLOS Comput. Biol. 12, e1004842 (2016).

27. Tennessen, J. A. et al. Evolution and Functional Impact of Rare Coding Variation from Deep Sequencing of Human Exomes. Science 337, 64–69 (2012).

28. Sharp, R. R. & Barrett, J. C. The environmental genome project: ethical, legal, and social implications. Environ. Health Perspect. 108, 279–281 (2000).

